# Fecal microbiota as a non-invasive biomarker to predict the tissue iron accumulation in intestine epithelial cells and liver

**DOI:** 10.1101/612168

**Authors:** Bingdong Liu, Xiaohan Pan, Liheng Yao, Shujie Chen, Zhihong Liu, Mulan Han, Yulong Yin, Guohuan Xu, Dan Wan, Xiaoshuang Dai, Jia Sun, Jiyang Pan, Huabing Zhang, Wei Wang, Li Liu, Liwei Xie

## Abstract

Iron is an essential trace mineral for the growth, systemic metabolism, and immune response. Imbalance of tissue iron absorption and storage leads to various diseases. The excessive iron accumulation is associated with inflammation and cancer while iron deficiency leads to growth retardation. Studies investigated in Kenyan infants and school children suggests that both low and high iron intake result in dysbiosis of gut microbiota. This would lead to the disruption of microbial diversity, an increase of pathogen abundance and the induction of intestinal inflammation. Despite this progress, in-depth studies investigating the relationship between iron availability and gut microbiota is not completely explored. In the current study, we established a murine model to study the connection between iron and microbiota by feeding mice with either iron-deprived or -fortified diet. To identify key microbiota related to iron levels, we combined the 16S rRNA amplicon sequencing with the innovated bioinformatic algorithms, such as RDA, co-occurrence, and machine learning to identify key microbiota. Manipulation of iron levels in the diet leads to systemic iron dysregulation and dysbiosis of gut microbiota. The bioinformatic algorithms used here detect five key bacteria that correlate with systemic iron levels. Leveraging on these key microbiotas, we also established a prediction model which could precisely distinguish the individual under either iron-deprived or iron-fortified physiological condition to further prove the link between microbiota and systemic iron homeostasis. This innovated and non-invasive approach could be potentially used for the early diagnosis and therapy of iron-dysregulation related diseases, e.g. anemia, inflammatory disease, fibrosis, and cancers.

## Background

Gut microbes are associated with host health (1). The dysbiosis disturbs the host systemic immune and metabolism balance, leading to the disease development and progression(2, 3). Although gut microbiota composition between twins is associated with heritage components, environmental factors significantly perturb the microbial homeostasis upon exposed to the different growth environments (4). Among these factors, dietary pattern shapes the gut bacterial microenvironment and further influences the microbial diversity. Most importantly, the diet composition contributes significantly to the gut flora homeostasis. For example, the plant- or animal-based diet leads to dramatic difference on the composition and diversity of gut microbiota in each population. Screening the gut bacteria diversity among 60 mammalians according to their dietary patterns indicated that the highest gut flora diversity in herbivores is followed by omnivores, and then carnivores (5). Studies have suggested that vegetarian-based population has a lower risk to develop chronic diseases including obesity, and diabetes than the western-diet-based populations (6–9). A recent investigation regarding the Mediterranean diet (MD) on gut microbiota suggested that MD is preferential to decrease the ratio of Firmicutes/Bacteroides, to increase the butyrate-producing bacteria in the gut, and to prevent the development of the low-grade inflammation (10).

The major composition of diet is protein, carbohydrate, and fat. Both rats and mice on high sucralose diets for 6 to 12 months leads to a lower proportion of Bacteroides, Clostridia, Bifidobacterial, Lactobacilli and total aerobic bacteria, followed with the induction of pro-inflammatory gene expression in intestinal epithelium (11, 12). Besides the major components, vitamin and mineral also play a pivotal role in shaping host health (13). As one of the essential trace elements, iron supports growth, oxygen transport, and metabolism. Both inadequate and excessive amount of iron intake are harmful to host. Except a small portion being absorbed by small intestine, the rest of unabsorbed iron passes into colon for bacteria, since iron is essential for most bacterial growth and survival (14–16). Thus, iron availability may impact the ecosystem of gut microbiota, further modulating the host health and metabolism. Several studies pointed out iron fortification may cause adverse effect on gut bacteria, especially leading to low grade inflammation to host. One study investigating the fortification of iron on infants in developing countries resulted in an adverse effect on gut microbiota and induction of gut pathogens and inflammation (17). In an iron fortification study among schoolchildren in Cote d’Ivoire, iron led to an induction of Enterobacteria, resulting in the low grade inflammation and a decrease of Lactobacilli (15).

Aforementioned, among these studies of the iron availability for gut microbiota, the phenotypical observation was made on the impact of iron towards the gut microbiota regarding the gut epithelial cell junction, host immune function and systemic inflammation. However, systemic understanding the role of iron on gut microbiota, especially utilizing the high-throughput sequencing and innovated bioinformatic analysis remains unexplored, especially the utilization of machine learning model to identify the key microbiota as the biomarker in predictive models. In the current investigation, we showed that gut taxa composition, and metabolic functions are perturbed by alteration of iron content in diet. This study is the first-time to combine high-throughput amplicon sequencing and bioinformatic analysis especially the machine learning to systemically explore the biological effect of iron on gut microbiota, in turn to perturb the host health and systemic metabolism.

## Methods

### Animals and diets

*C57BL/6J* mice were raised at SPF animal facility of Guangdong Institute of Microbiology (GDIM) in a 12/12 dark-light cycle with *ad lithium* free access to food and water. The animal protocol was proved by the Institute Animal Care Use Committees of GDIM (Permission #: GT-IACUC201704071).

*C57BL/6J* mice at 6-week old of age were fed with Ain93G-based diet supplemented with different amount of iron, Control (∼33 ppm), low-iron (<3 ppm) or high-iron (∼200 ppm) for 12-week. Each group had 9 mice, housed in the same cage. The formula was listed in Supplementary Table 1. Each mouse was weighted every week to monitor the body-weight. Specialized AIN93G-based diets were purchased from Changzhou SYSE Bio-Tech Co. LTD.

### Blood parameter measurement

The tail was sterilized with 75% ethanol pad before blood was collected from tail-vein to measure the hemoglobin and hematocrit.

### Feces collection

Fresh feces were collected with a sterilized tube and frozen right away in −80°C freezer for further analysis.

### Iron assay

Fecal iron content was measured following the standard ferrozine iron assay protocol as described before (18).

### RNA isolation and reverse transcription and Quantitative Real-time PCR (qRT-PCR)

Total RNA was isolated and reverse transcribed to cDNA, followed with the qRT-PCR as described before (19).

### 16S rRNA amplicon sequencing

Total fecal bacterial DNA was isolated with the Mobio PowerSoil® DNA Extraction kit. DNA concentration was determined by BioSpec-nano (Shimadzu Corporation, Japan). 50 ng DNA was used for PCR-amplification with Q5 High-Fidelity DNA Polymerase (NEB) targeting to the V3-V4 region of bacterial 16S rRNA gene, followed with purification with AMPure XP (Beckman). The PCR product was indexed with Illumina sequencing primer-set by utilizing the Phusion HF Taq Polymerase (NEB), followed with gel purification. DNA concentration was quantified with Qubit-3 (Thermo Fisher). The PCR product was sequenced on an Illumina HiSeq2500 platform. Primers for library construction were listed in Supplementary Table 1.

### Bioinformatic and statistical analysis

After sequencing, paired-end raw reads were collected from BioMarker (BMK). For each pair-end sequencing file, the low-quality reads, adaptor, and barcode were trimmed using the FASTX-Tool kit and the pair-end sequencing files were merged. All fastq files were quality-controlled with QIIME 1.9.1 workflow (20). A default similarity threshold at 97% was defined. The BIOM file was used for downstream analysis with QIIME1.9.1 pipeline and R.

**α- and β-diversity** For α-diversity, Observed_OTUs, Shannon and Simpson were rarified at the same sequencing depth were used to present the observed species and diversity of the gut microbiota. β-diversity was assessed using the Binary_jaccard, Bray_curtis, Weighted_unifrac and Unweighted_unifrac to calculate the distance between groups. Principal Coordinates Analysis (PCoA) were plotted by ‘ggplot2’ (21) and ‘vegan’ (22) of R. The relative abundance of microbial structure was plotted by ‘pheatmap’ package of R. **BugBase** was performed by utilizing BugBase algorithm with integration of KEGG (23). **Analysis of Similarity (ANOSIM)** was performed with “vegan” package of R, based on Bray Curtis distance. **Linear discriminate analysis effect size (LEfSe)** was performed to identify the difference of microbial structure between groups with the default parameter (24). **PICRUSt** was used to explore the difference of metabolic pathway between each group against the KEGG (25). **Redundancy analysis (RDA)** was performed by using ‘vegan’ and ‘ggplot2’ package of R (21, 22). The length of arrow represented the constraint effect of different environmental variables exerted on the sample distribution in two-dimension. **Co-occurrence analysis** was performed by ‘igraph’ package of R. All nodes were annotated by different color on phylum level. Communities of three network were determined by the fast greedy modularity optimization algorithm (26). Circle bar was plot according to eigenvector centrality scores to estimate the importance and betweenness of each node (27). The index of radar plot estimated by the graph theory algorithm was plotted by ‘fmsb’ package of R. **Random forest** was performed with ‘randomForest’ package of R (28). **Least absolute shrinkage and selection operator (LASSO)** was performed by ‘glmnet’ package of R (29).

### Data availability

The raw sequencing file for all samples have been deposited in NCBI under the Bioproject: PRJNA506862.

## Results

### Dietary manipulation of iron levels perturbs systemic iron homeostasis

In an attempt to manipulate the systemic iron dysregulation to further understand the precise iron-mediated regulatory mechanisms on gut microbiota, 6-week old *C57BL/6J* mice were fed with an AIN93G-based diet for 12-week to induce the iron-deficiency or iron-overload (Fig. 1A). The results showed that iron-deprived diet led to the growth retardation, lower hemoglobin and hematocrit (Fig. 1 B, C, D). We also observed that iron levels were significantly reduced in epithelial cells of duodenum (SI), liver, BAT and iWAT, while FeE diet led to iron-overload only in SI and liver but not in BAT and iWAT, compared with the mice with AIN9G diet (Fig.1E). The unabsorbed iron in the diet was passed into colon and packed into the feces as shown in fecal iron content (Fig. 1E). The expression of iron transport-related genes was differentially expressed, such as *Slc11a2, Slc40a1, Slc39a14* and *TfRc* in duodenum and *TfRc* and *Slc39a14* in liver (Fig. 1F).

**Figure 1.**
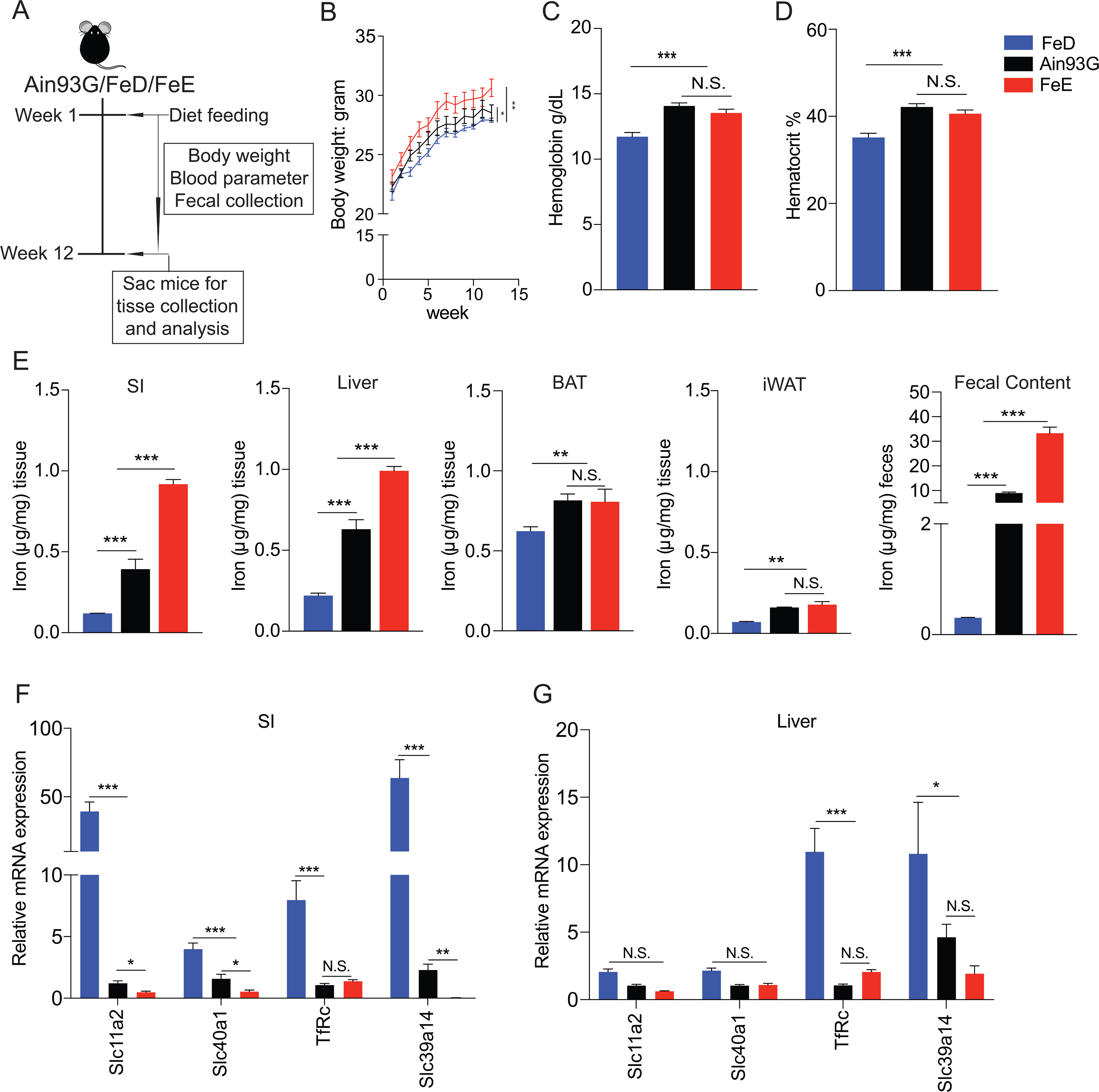
Different dietary iron level in diets perturbs systemic iron homeostasis and alters the iron-related parameter levels. *C57BL/6J* mice were fed with AIN93G-based diets including iron-deficiency (FeD), iron-fortified (FeE) and AIN93G (Control) for 12 weeks to disrupt systemic iron homeostasis. (A) Experimental scheme to access the role of iron on fecal microbiota. (B) Body weight curve of *C57BL/6J* mice under three different diets. (C-D) Hemoglobin and hematocrit for *C57BL/6J* mice under three different diets. (E) Tissues (small intestine/SI, liver, brown adipocyte/BAT, inguinal white adipocyte/iWAT) and fecal iron levels for C57BL/6J mice under three different diets. (F-G) Iron transport-related gene 是 expression (*Slc11a2, Slc40a1, TfRc, Slc39a14*) in epithelial cells of duodenum and liver. Each bar or point represents the average data from 9 mice. * *p*<0.05, \*\**p*<0.01, \*\*\**p*<0.001, N.S.: not significant.

### Iron-dysregulation significantly perturbs the gut microbiota diversity and metagenomic function

After 12-week feeding, 27 fecal samples were collected and sequenced, generating 935282 high quality ∼420-bp paired-end reads with average 34640 reads for each sample. These high-quality reads were used to calculate the relative abundance against the Greengenes database. To determine whether sequencing depth for each sample was enough to capture the diversity of gut microbiota, random sampling was performed to plot the species accumulation curves (SAC). The SAC showed that with the increased sampling, OTUs reached the asymptote (Fig. S1A). The Good’s coverage curves demonstrated that all samples reached saturation within 5000 paired-end reads. The sequencing coverages were near 100%, suggesting that the sequencing depth was sufficient to capture the diversity of gut microbiota for all groups (Fig. S1B, S1C). Rarefaction analysis based on α-diversity clustering showed that different iron level in diet significantly perturbed the number of species and diversity between low-iron (FeD) and high-iron (FeE) diet feeding, represented by Observed_OTUs (*p*=0.006), Shannon (*p*=0.014) and Chao1 (*p*=0.0022) index, respectively (Fig. 2A). To display the microbial variability between groups, β-diversity analysis was calculated with the Bray_curtis, Binary_jaccard, Unweight_unifrac and Weighted_unifrac algorithms. The space distance among groups presented a symmetrically different distribution (Fig. S2A-D). It was showed that on PCoA1 axis, space index was significantly different between any two groups (Fig. 2B). By comparing the relative abundance on species level among three groups, Kruskal non-parameter test (*p*<0.05) showed that different iron content in diet significantly altered the gut microbiota composition (Fig. 2C). Moreover, LEfSe analysis identified 34 (AIN93G), 18 (FeD), and 11 (FeE) bacterial taxa that were differentially altered with LDA score >2, showing that there was significant structural difference among three groups (Fig. 2D). All these results demonstrated that iron may be a key modulator towards the gut microbiota diversity.

**Figure 2.**
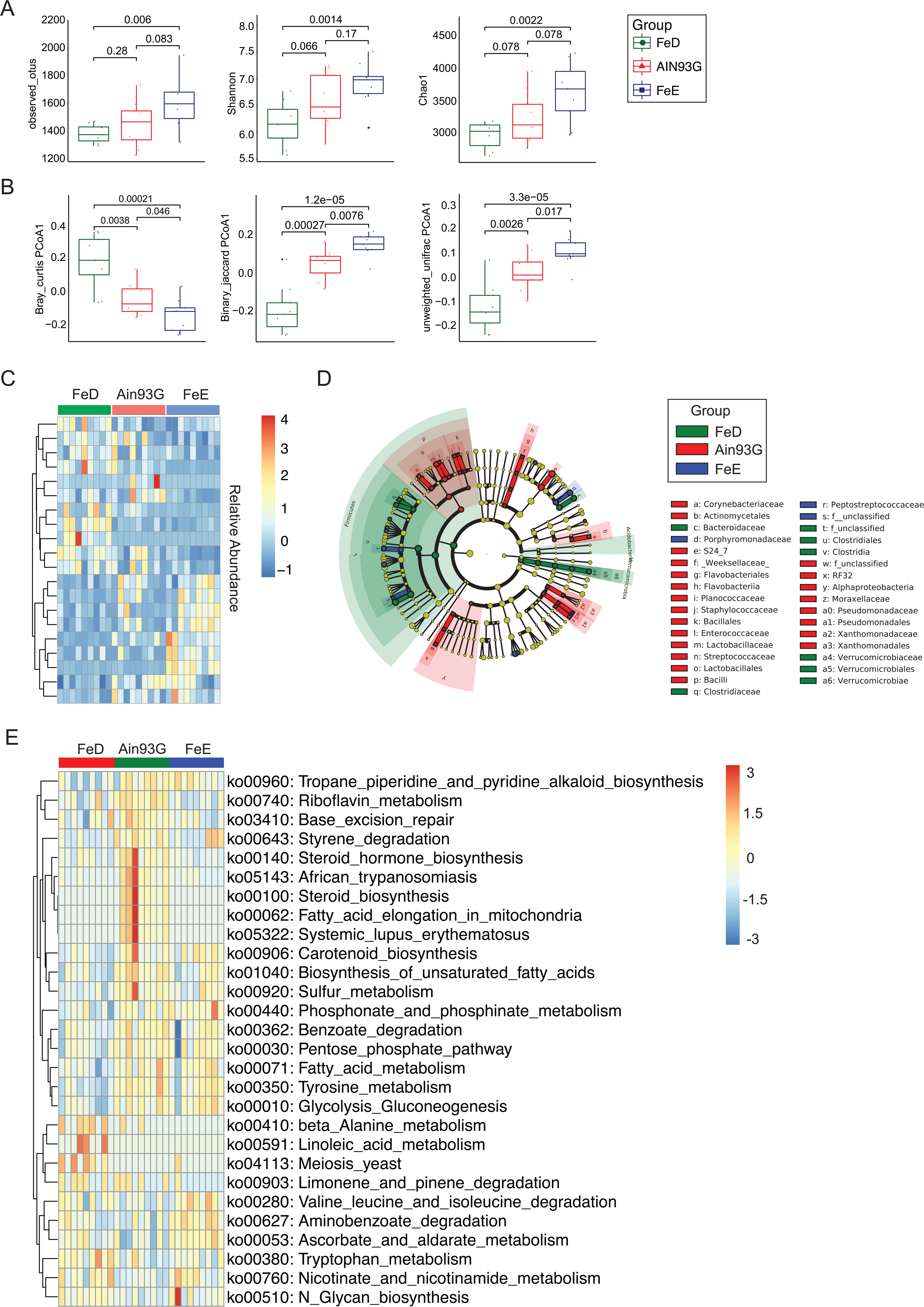
Iron-dysregulation significantly perturbs the gut microbiota diversity and metagenomic function. Feces were collected from *C57BL/6J* mice under three different diets and its bacterial DNA were isolated. V3-V4 region of 16S amplicon was sequenced for bioinformatic analysis to assess the microbial composition. (A) Box plots of α-diversity index (Observed_otus, Shannon, and Chao1). (B) Box plots of β-diversity index of PCoA1 represented by the space distance among groups (Bray_curtise, Binary_jaccard, Unweighted_unifrac). (C) Heatmap for relative abundances of significant difference at species level among three groups (Kruskal non-parameter test (*p*<0.05)). (D) Linear Discriminant Analysis (LDA) Effect Size (LEfSe) plot revealed the taxonomic biomarkers associated with the dietary iron levels. The threshold for LDA score was 2 (*p*<0.05). (E) To predict the metagenome function, heatmap of PICRUSt analysis showed significant KEGG pathway among three groups.

In order to understand whether the alteration of gut microbial diversity can cause functional changes, BugBase was performed. However, the relative abundance of gut microbiota under different phenotypes were not significantly different (Fig. S3). To predict the metagenomic function, PICRUSt was used to assess the functional difference between microbiota among three groups. It was shown that fatty acid metabolism, tyrosine metabolism and gluconeogenesis were slightly more abundant in gut taxa of mice under diet with high iron content. The linoleic acid metabolism, β-analine metabolism and limogene and pinene degradation were enriched in bacterial taxa under low iron condition (Fig. 2E). These results confirmed that the functional and phenotypical pathways were altered, following the gut microbiota diversity change.

### Gut microbiota alteration strongly associated with dietary iron content

The specialized diets have significantly perturbed mice physiological parameters. Whether these changes were related with gut microbiota remains unknown. Thus, we performed RDA analysis to further reveal the relationship between the iron-based physiological parameter and gut microbiota. This analysis indicated that 26.72% of the variance could be interpreted by six environmental factors (Supplementary Table 2), which confirmed that systemic iron levels could significantly alter the population of the gut microbiota at phylum level and samples from three groups were obviously separated (Fig. 3). According to the Monte Carlo permutation test, physiological parameters, such as Hemoglobin (*p*=0.004), Hematocrit (*p*=0.004), SI iron content (*p*=0.006) and fecal iron content (*p*= 0.023) played an essential role in clustering the distribution of bacterial taxa in three groups (Fig. 3). To further confirm the RDA analysis, ANOSIM based on the Bray Curtis distance also indicated a statistical difference between three treatments with a statistic test (*R=*0.537 for *p*<0.0001) (Fig. S2E). Both RDA and ANOISM analysis proved that dietary iron level plays an essential role in separation and clustering of the gut microbiota in each group.

**Figure 3.**
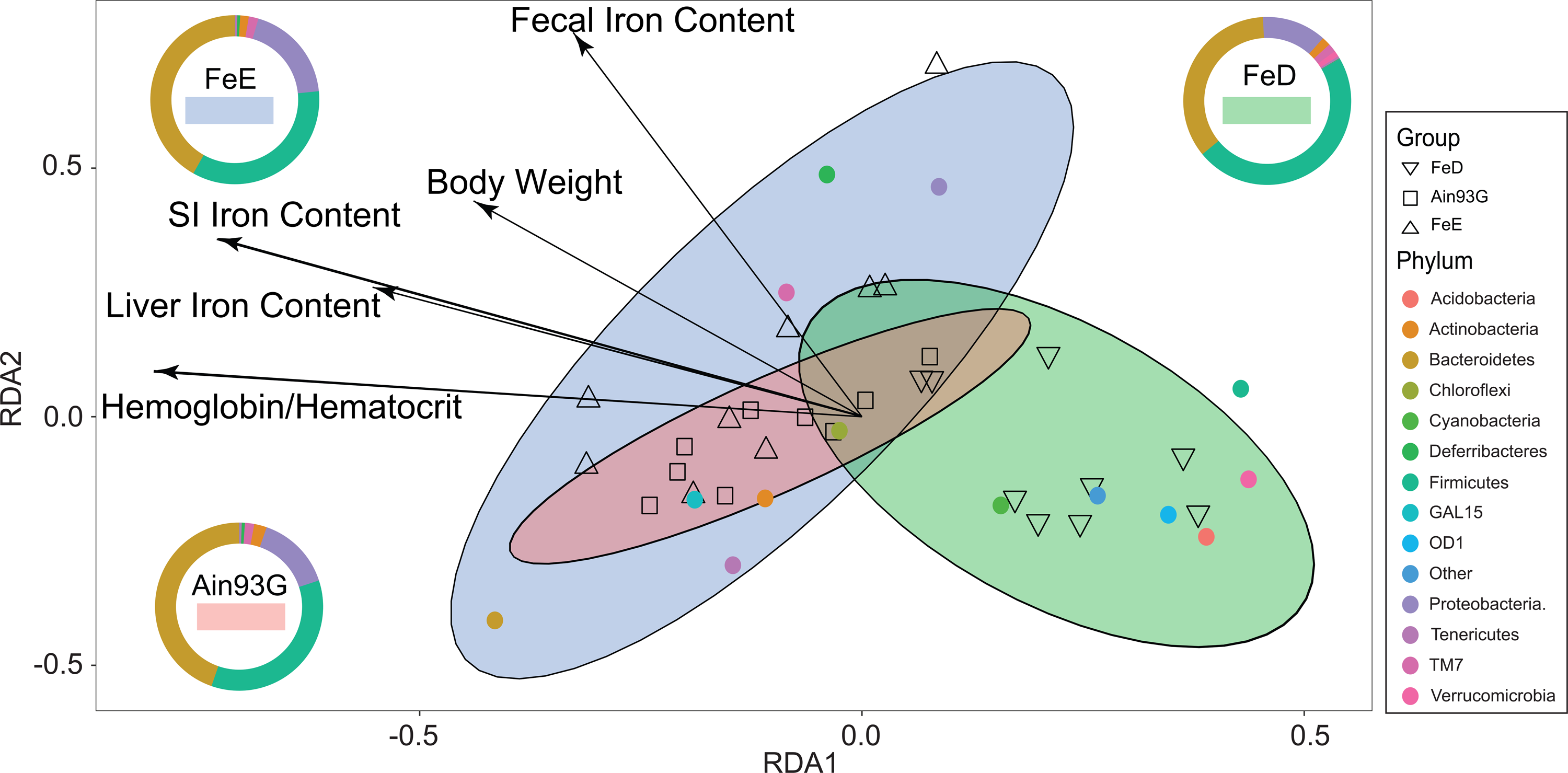
Redundancy analysis (RDA) bi-plot of bacterial diversity and iron-related physiological parameter. RDA was performed on the relative abundance obtained from the 16S rRNA amplicon sequencing and environmental factors (body weight, hemoglobin, hematocrit, SI iron content, liver iron content and fecal iron content).

### Iron supplementation perturbs the gut microbial community network

To determine whether iron is able to influence the gut microbiota community network, the graph theory algorithm and co-occurrence analysis were performed. In order to systemically understand the interaction between bacterial taxa, the graph theory including the transitivity, graph density, degree centralization, number of vertices and number of edges on a radar plot indicated that the iron content in diet did not significantly change the systemic complexity of gut bacteria (Fig. S4 and Supplementary Table 2). Thus, to further explore the bacterial interaction in response to the systemic iron level change, we performed the co-occurrence analysis through the collection of the prevalence of bacterial species from each group in terms of relative abundance at species level as detailed in Supplementary Table 2. The plotted taxa were the relative abundance present in at least 70% of the samples in each group and were used for co-occurrence analysis (Supplementary Table 2). The statistical analysis through spearman correlation, species plotted for co-occurrence network separately suggested that the microbial community among three groups were significantly different (*r*>0.6, *p*<0.05, Supplementary Table 3). By applying the greedy clustering method, FeD group was sub-divided into 6 sub-communities, while AIN93G and FeE groups had 5 sub-communities (Fig. 4 and Fig. S5). Within this network, iron indeed reorganized the network, providing a clue that a threshold for iron levels in diet may present. This threshold may be a determinant factor affecting the cluster of the sub-community as shown in diversity analysis.

**Figure 4.**
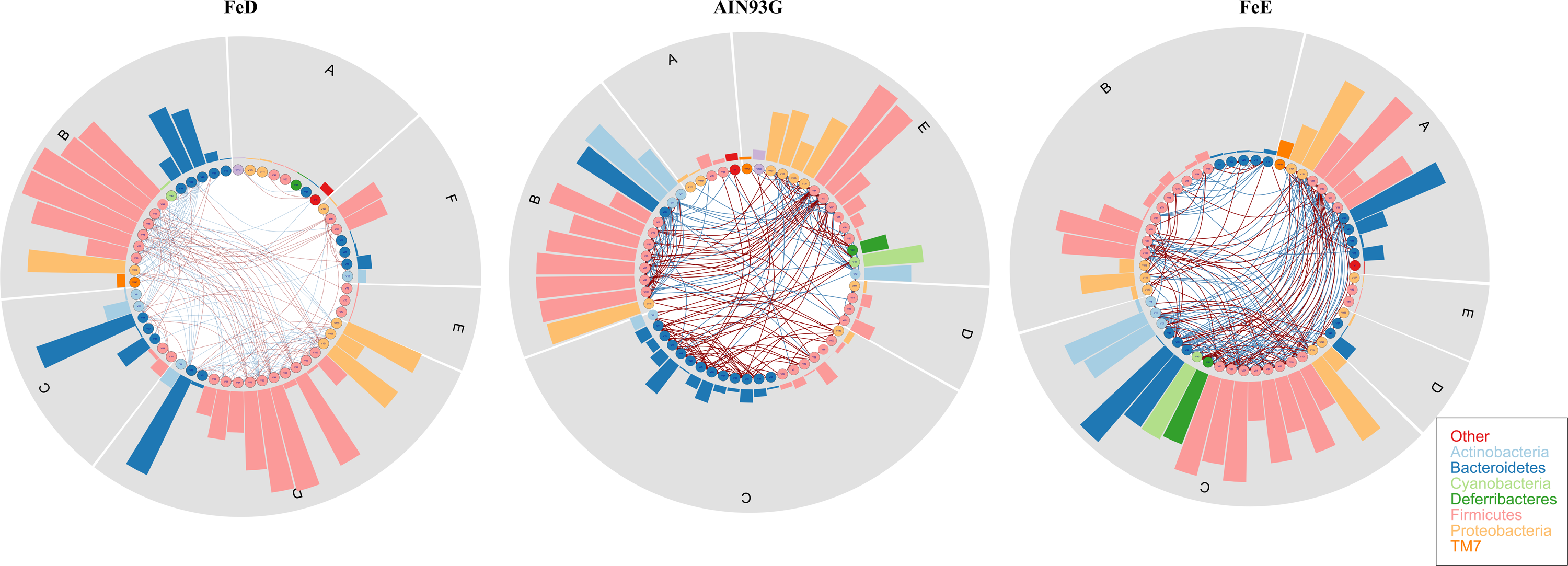
Co-occurrence network analysis and prevalence of bacterial species among different dietary treatment. Network plot describes co-occurrence of bacterial species among all samples. (A) FeD, (B) AIN93G, (C) FeE. The blue and red line indicates the negative and positive interaction between bacterial taxa, respectively. The number of segments of each circle (gray color) revealed the number of sub-communities of each group. The height of the bar in each segment represented the significance of the corresponding bacteria. The bacterial species in co-occurrence analysis were only those relative abundance present in at least 70% of the samples in each group.

### Identification of the key bacteria associated with the systemic iron dysregulation by random forest

We have shown that iron regulates host physiology and metabolic balance and also significantly influences the structure of microbial community. This resulted in 120 bacterial abundance significantly changed by the iron level in diets (kruskal non-parameter test with *p*<0.05) made it difficult to distinguish their importance. Here, we introduced the random forest algorithm to identify the valuable microbiota among three groups in response to the systemic iron dysregulation. By applying the five-fold cross-validation on a random forest model among total 27 samples in the discovery phase on species level, ∼70 million decision trees from ten trials were generated, resulting in 4 optimal species markers selected with consideration of the lowest mean error rate and standard deviation (Fig. 5A). Moreover, through ten trials of analysis, the top five taxa were consistent (Fig. S6A). Thus, we selected the top five candidates as the key microbiota in response to the iron-mediated systemic dysregulation (Fig. 5B). This included g_Parabacteroides (V19), f_Pepostreptococcaceae (V83), s_Akk_muciniphila (V143), s_perfingens (V70) and o_clostridiales (V63) (Fig. S7A-B). Furthermore, the partial dependence plot showed that the iron is involved in the classification of these five key microbiotas, based on their relative abundance (Fig. S7C). These five key taxa were highly correlated with the physiological parameters (Fig.5C and Fig. S7D). Given the key position of five microbiotas among the community in response to the systemic iron dysregulation, based on their relative abundance, we built a prediction model, which was able to recognize the target samples. ROC curves suggested that samples could be well distinguished with AUC value of 99.4%, 88.9% and 100%, respectively (Fig. 5D-F, S6B).

**Figure 5.**
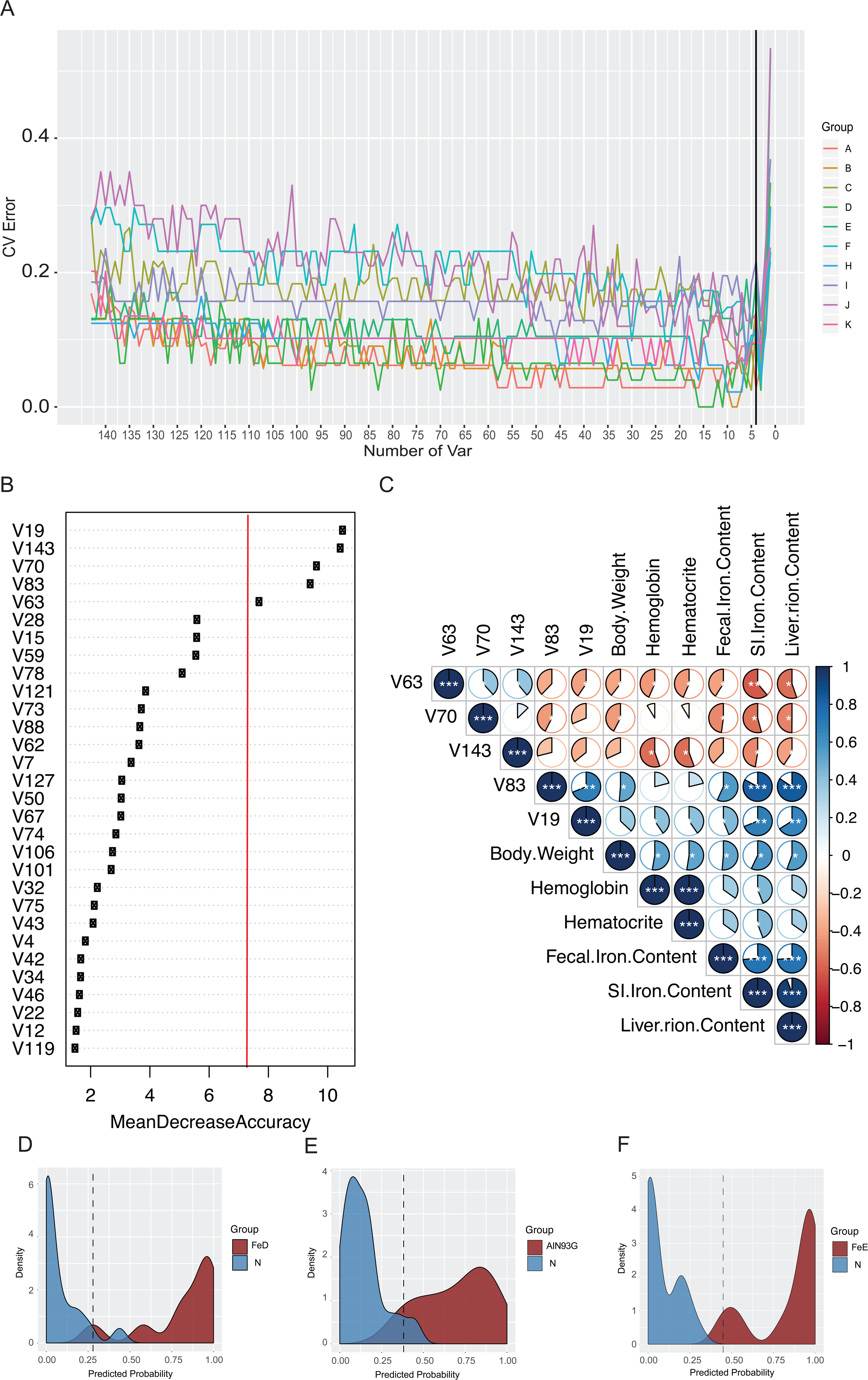
Identification of the signature gut microbiota associated with the systemic iron dysregulation by random forest. To detect signature bacterial marker in response to the systemic iron dysregulation, we performed five-fold cross-validation on a random forest model among 27 samples (9 for each groups) in the discovery set. (A) 4 bacterial markers at species level were selected as the key biomarker by random forest. Red line illustrates the number of key bacteria in the discovery set. (B) The relative abundance of each bacteria in the predictive model was assessed using the Mean Decrease Accuracy (MDA) in 27 datasets. The red line illustrates five key bacteria filtered by random forest via the fivefold cross-validation. (C) The corplot analysis plots the correlation between five signature bacteria and environmental factors (Body weight, Hemoglobin, Hematocrit, Fecal Iron Content, SI Iron Content, and Liver Iron Content). (D-F) The mathematic model based on the 5 key bacteria was used to predict the sampling group. The probability plot indicates the prediction probability. The dark red color indicates the predicted group and the blue color is the non-predicted samples. Black dashed line represents the lowest probability to predict the samples belonging to the correct group.

### Relative abundance of gut microbiota-based prediction on the systemic iron status

Random forest classifier identified 5 key microbiotas in response to systemic iron homeostasis with tight correlation between five key microbiota and iron-related physiological parameters. Thus, we speculated whether the relative abundance of gut microbiota could be used as a biomarker to predict tissue iron content. Here, we constructed LASSO regression models, which could help us select the best independent parameter. With the λ selected from the LASSO regression models, we identified the essential microbial taxa to build the prediction model for each environmental factor (Fig. S9, Fig. 6A, and 6D, Fig. S8, Fig. S10A, and 10C). Utilizing these models to predict SI, liver, fecal iron content and hemoglobin resulted in *R*^*2*^ of 99.7%, 96%, 65% and 67%, respectively (Fig. 6B and 6E, Fig. S10B and 10D), especially the SI and liver iron contents associated with high prediction confidential. Leveraging on two machine learning models, including random forest and LASSO regression model, key microbiota from two models were merge to be used as key microbiota (4 and 2 taxa biomarkers for SI and liver) to predict the SI and liver iron content (Fig. 6C and 6F).

**Figure 6.**
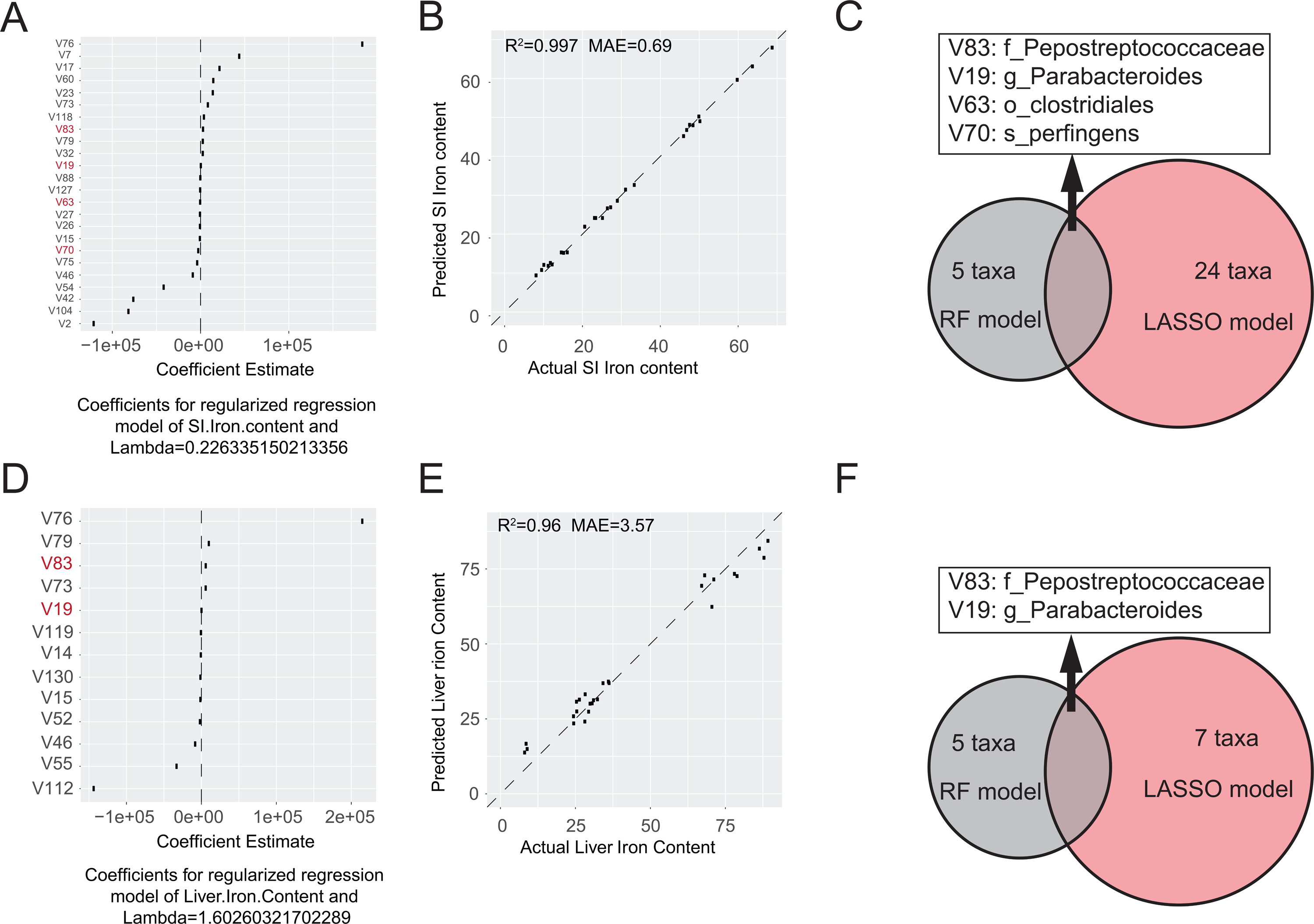
LASSO regression model to utilize the relative abundance of bacterial taxa to predict systemic iron levels. SI (A) and Liver (D) iron content-associated taxa were selected from LASSO regression model (L1-regularization) with optimized λ with repeated 6-trial. Taxa from LASSO regression model to establish mathematic model could utilize the relative abundance of gut microbiota to predict SI (B) and Liver (E) iron content with R^2^ of 0.997 and 0.96, respectively. The overlapping taxa between two machine learning model to generate the key bacterial taxa that were used to predict SI (C) and liver (F) iron content.

## Discussion

Iron plays a fundamental role in various aspect of life, such as oxygen transport, metabolism, and immune defense. Small intestinal absorption accounts for 2 to 3 mg/day for human and the rest of iron passes into the colon for gut microbiota (14). However, whether fecal and systemic iron level influence the homeostasis of gut microbiota remains unknown. To precisely understand the impact of iron on gut microbiota, fecal samples from *C57BL/6J* mice were collected and its microbial diversity and metagenomic function were assessed. From the current investigation, we have shown that the iron levels in diet is a key factor contributing significantly to the gut microbiota sub-community formation. Most importantly, based on the machine learning algorithm, we built the prediction models, which help us identify the key microbiota in response to the systemic iron challenge, and establish the connection between key microbiotas and tissue iron content. By utilizing these newly identified biomarkers, we could accurately predict SI and liver iron content with *R*^*2*^ of 99.7% and 99.6%, respectively.

Iron is also a critical mineral for gut microbiota. Studies from multiple approaches demonstrated the iron availability directly impacts the bacterial ecology under both *in vivo* and *in vitro* conditions. In human studies, the availability of iron may present different effect on the diversity of gut flora. Infants at their first three months had different response to the milk with different iron fortification when compared with the breast milk. For the new born infants, iron-fortification is required for gut flora colonization from the standardized prepared cow’s milk (30). However, the follow-up study from the same group showed that in the later period of milk feeding, low iron-fortification in the cow’s milk manipulated similar observation of flora type as that in breast milk fed infants in the first three months (31). In animal studies, swiss-Webster mice fed on iron-deprived diet resulted in elevation of the anaerobes in the colon (32). Individual on iron deficiency followed with iron supplementation also significantly modified the bacterial diversity and metabolites levels, especially the butyrate and propionate levels (33). These human and animal studies have established the connection between iron concentration and overall gut microbiota composition and complexity, but these all based on the microbial sampling, phenotypic observation and the relative abundance, which still lacks of the precise evaluation and prediction approach to dissect the link between gut microbiota and host physiology.

Currently, the widely used approaches include α-diversity, β-diversity, and LEfSe to calculate the difference between groups. These approaches are straightforward to identify the significant difference, while it may result in false-positive conclusion and also lack of the cross-validation. Recently, in a randomized controlled study conducted in Swedish healthy infants at 6 months old of age with different ferrous iron administration in milk, LEfSe algorithm is used to identify bacterial taxa associated with different iron content in formula. It was shown that high-iron formula is associated with lower relative abundance of Bifidobacterium and Lactobacilum (34). In our attempt, the relative abundance of Lactobacilum decreases gradually when additional iron is supplemented in diet. However, the relative abundance of Bifidobacterium was not changed among groups. In our study, other than comparing the relative abundance among groups, we incorporated multiple approaches to understand the link between systemic iron homeostasis and the dysbiosis of gut microbiota, such as RDA, co-occurrence, and machine learning model to determine the overall impact of dietary iron on microbial composition, sub-community, interaction, and clustering. Among them, RDA leverages on the environmental factors and relative abundance of gut microbiota to determine how environmental factors contribute to the clustering of bacterial taxa and sub-community formation. Here, iron is a major factor to influence the iron-related parameters and associated with the tissue iron levels, which contributes to the clustering of gut microbiota in each group. This was represented by both RDA plot and co-occurrence via the bacterial interaction. The machine learning approach is introduced in microbial studies to identify the signature microbiota, which were utilized as diagnosis markers to predict the disease progression, such as irritable bowel syndrome (IBS) (35), breast cancer (36), hepatocarcinoma (37). In a gut microbial study of IBS patients, relying on the machine learning and 5-fold cross-validation to reduce the complexity of sequencing data. The gut microbial signature could be used to discriminate the healthy individuals and patients with different degree of symptom with LASSO regression model (35). Furthermore, gut microbial signature is also used in combination with the cross-validation and random forest to identify potential diagnosis microbial biomarkers for early diagnosis of hepatocarcinoma and breast cancer (36, 37). Obviously, the robust statistical approaches could overcome weakness of just the difference on relative abundance to identify the key microbiota. In our study, 5-fold cross-validation with random forest identified five key microbiotas, which highly correlated with the environmental factors. LASSO regression model further confirmed the essential connection between the identified microbiota and the tissue iron with a high confidential rate.

Furthermore, cancer development is associated with the microorganism in some organs, for example, *helicobacter_pylori* and gastric cancer, or *papillomavirus* and cervical cancer. Recent studies have shown that gut microbiota contribute to the pathogenesis and progression of colorectal cancer. Some bacterial species such as *Clostridium_septicum, Enterococcus_faecalis, Streptococcus_bovis, Bacteroides_fragilis, He licobacter_pylori, Escherichia_coli and Fusobacterium spp* may participate in the development and progression of CRC (38, 39). Clostridium_septicum as a virulent pathogen has been shown to involve in disease development, e.g. colorectal malignancy, and hepatic carcinoma, but this may happen with involvement of other clostridial species such as Clostridium_Chauvoei, Clostridium_Novyi, and Clostridium_Perfringens (40). Clostridium_septicum infection and severity may be associated with the dysfunction of neutrophil. Neutrophils-mediated immune system plays a fundamental role in restrict colorectal tumor development. A recent study indicated that neutrophils deletion leads to the sporadic colon tumorigenesis with increased bacterial growth in tumor, proliferation of tumor cell, and inflammatory response mediated by interleukin 17 (41). 16S rRNA-sequencing data suggested that dysbiosis of gut microbiota, especially the Clostridiales, and Akk makes significant contribution to the development and progression of tumor in neutrophils-deficient mice (41). Clostridium_perfringens is another member of Clostridiaceae family in human and animal intestine and is associated with various systemic and enteric diseases such as diarrhea, colitis, and colon cancer (42). Here, we identified iron deficiency environment may be suitable for Clostridiacae family growth, while Peptostreptococcaceae family is enriched in intestine of mice fed with iron-fortified diet. Peptostreptococcaceae appears to be over-represented in the gut of both colorectal cancer patients and animal models (42). This family of bacteria secretes more than 20 identified toxins or enzymes that could potentially impact the homeostasis of gut microbiota and induce the pathogenesis in the host gut, leading to colitis and tumor development (42).

Cancer development and progression is a complicated cascade with multiple factors involved, such as nutrients, genetics, and bacteria. Iron dysregulation is associated with the development of various types of inflammatory diseases and cancers (43). Furthermore, the meta-analysis of the epidemiological evidence supported the positive association between dietary iron intake and higher cancer risk (44). In various cancers, such as liver, breast, and colorectal cancer (45, 46), it has been shown that the development is primarily followed with abnormal iron absorption and progressed with the accumulation of excessive amount of free iron. Thus, development of a prediction model for the early diagnosis is necessary. Here, in our study, we present a mathematic model that establish a link between gut microbiota and tissue iron level. The application of the current study is to utilize the relative abundance of gut microbiota to predict tissue iron level as a biomarker to monitor the iron accumulation related diseases development and progression. In present stage, our findings provide evidence support the link between systemic iron and gut microbiota and the prediction model could be potentially used to predict the SI and liver iron levels for the iron-related disease early diagnosis (Fig. S10).

## Supporting information

Sequencing depth was assessed via the random sampling

???-diversity and ANOSIM analysis demonstrate dietary iron manipulation perturbs the gut microbiota diversity

Metagenomic prediction of microbial phenotypes using Bugbase

Dietary iron content does not change the complexity of gut microbiota

Co-occurrence network analysis and prevalence of bacterial species among different dietary treatment

Identification of the signature taxa by random forest

Identification of the signature taxa by random forest

Selection of hyperparameter (???) for LASSO regression model.

LASSO regression model to utilize the relative abundance of bacterial taxa to predict systemic iron levels

Schematic representation of the link between gut microbiota and systemic iron homeostasis.

Diet Fomular

Analysis meta data

Cross-validation and random forest

## Conflict of Interest

The authors declare that there is no conflict of financial or research interest

## Author contribution

LW. X., GH. X., YL. Y. W. W. and L. L designed the experiments. BD. L. and ZH. L. performed data analysis. ML. H. and D.W. performed animal work and collected animal feces. XH. P, SJ. C., J. S. and LH. Y. performed the in vivo animal experiments. BD. L., LW.X., JY. P. and HB. Z. drafted the manuscript.

## Acknowledgement

This work was support by ‘GDAS’ Project of Science and Technology Development (Grant No. 2018GDASCX-0806) to Liwei Xie and the High-level Leading Talent Introduction Program of GDAS (Grant No. 2016GDASRC-0202) to Yulong Yin, National Natural Science Foundation of China (Grant No. 81670783) and, the Natural Science Foundation of Guangdong (Grant No. 2017A030313473) to Jia Sun.

## Supplementary Figure Legends

Fig. S1 **Sequencing depth was assessed via the random sampling**. Random sampling and goods_coverage were used to estimate whether the sequencing depth is enough to capture the diversity of gut microbiota. (A) species accumulation curves (SAC). (B-C) Goods_coverage plot.

Fig. S2 **β-diversity and ANOSIM analysis demonstrate dietary iron manipulation perturbs the gut microbiota diversity.** (A-D) β-diversity index: Binary_jaccard, Bray_curtis, Unweighted_unifrac, and Weighted_unifrac. (E) Analysis of similarity (ANOSIM) indicates a statistically significant difference between three treatments with a statistic test (*R=*0.537 for *p*<0.0001). Red dashed line illustrates the R value at 0.537.

Fig. S3 **Metagenomic prediction of microbial phenotypes using Bugbase**. Utilizing the 16S amplicon sequencing information, Bugbase algorithm was used to predict the organism-level coverage of functional pathways and phenotypes, such as gram positive or negative, aerobic or anaerobic, potential_pathogenic, and stree_tolerant.

Fig. S4 **Dietary iron content does not change the complexity of gut microbiota**. Radar plot with parameter of transitivity, number of edges, number of vertices, degree of centralization, and graph density demonstrates that microbial complexity in diet is not significantly changed by dietary iron level.

Fig. S5 **Co-occurrence network analysis and prevalence of bacterial species among different dietary treatment.** Network plot revealed the systemic interaction between each bacterium among diet with different dietary iron levels, (A) FeD, (B) AIN93G, (C) FeE.

Fig. S6 **Identification of the signature taxa by random forest**. To detect key bacterial marker, we conducted 5-fold cross-validation for 10 trials with seed number 1000 to generate 70 million decision trees. (A) The relative abundance of each bacteria taxa in the predictive model was assessed using the Mean Decrease Accuracy (MDA) in 27 datasets. Each graph represents one trial. (B) ROC curve for the training set. The AUC was 0.994, 0.889, and 1 for FeD, AIN93G and FeE.

Fig. S7 **Identification of the signature taxa by random forest**. (A) The detailed information of five key microbiota identified by random forest. (B) Bar plots show the relative abundance of each taxa classified by random forest. (C) The partial dependent probability plot illustrates the relationship between the relative abundance of each taxa identified by random forest and the probability to be classified under certain dietary iron level. (D) The corplot revealed in correlation between five core microbiota and environmental factors.

Fig. S8 **Selection of hyperparameter (λ) for LASSO regression model**. With the increasement of hyperparameter (λ), which exerts regression coefficient of all independent values in LASSO regression model approaching zero gradually, the number of independent values shrank. In each plot, the left line represents as a minimum λ that is the least MSE for the most accurate LASSO model, and the other one is chosen for the biggest λwithin one standard error of the minimum λ to construct the simplest LASSO model.

Fig. S9 **LASSO regression model to utilize the relative abundance of bacterial taxa to predict systemic iron levels**. Fecal iron content (A) and Hemoglobin (C) associated taxa were selected from LASSO regression model (L1-regularization) with optimized λ with repeated 6-trial. Taxa from LASSO regression model to establish the mathematic model could utilize the relative abundance of gut microbiota to predict Fecal iron content (B) and Hemoglobin (D) iron content with R2 of 0.65 and 0.67, respectively.

Fig. S10 **Schematic representation of the link between gut microbiota and systemic iron homeostasis**. Systemic dysregulation of iron metabolism is associated with various kinds of diseases development and progression, such as anemia, inflammatory diseases, and cancer. In our murine model, we combined the high-throughput sequencing and bioinformatic analysis, especially the machine learning approach dissect the link between gut bacteria and systemic iron homeostasis, providing a novel and alternative approach for early diagnosis of iron dysregulation-related diseases.

