## Supplementary figures and images for "Fecal microbiota as a non-invasive biomarker to predict the tissue iron accumulation in intestine epithelial cells and liver"

### ???-diversity and ANOSIM analysis demonstrate dietary iron manipulation perturbs the gut microbiota diversity

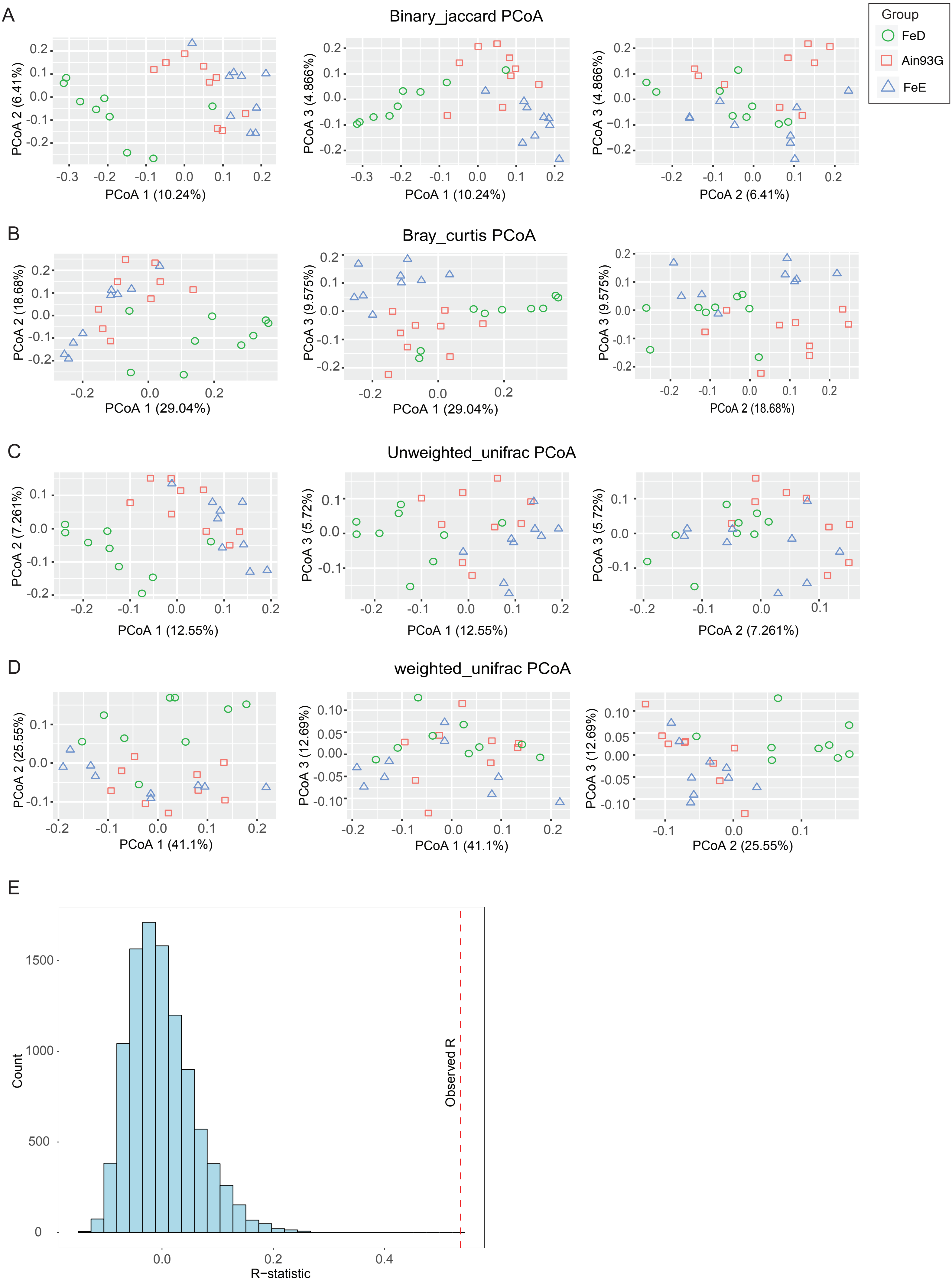

### Co-occurrence network analysis and prevalence of bacterial species among different dietary treatment

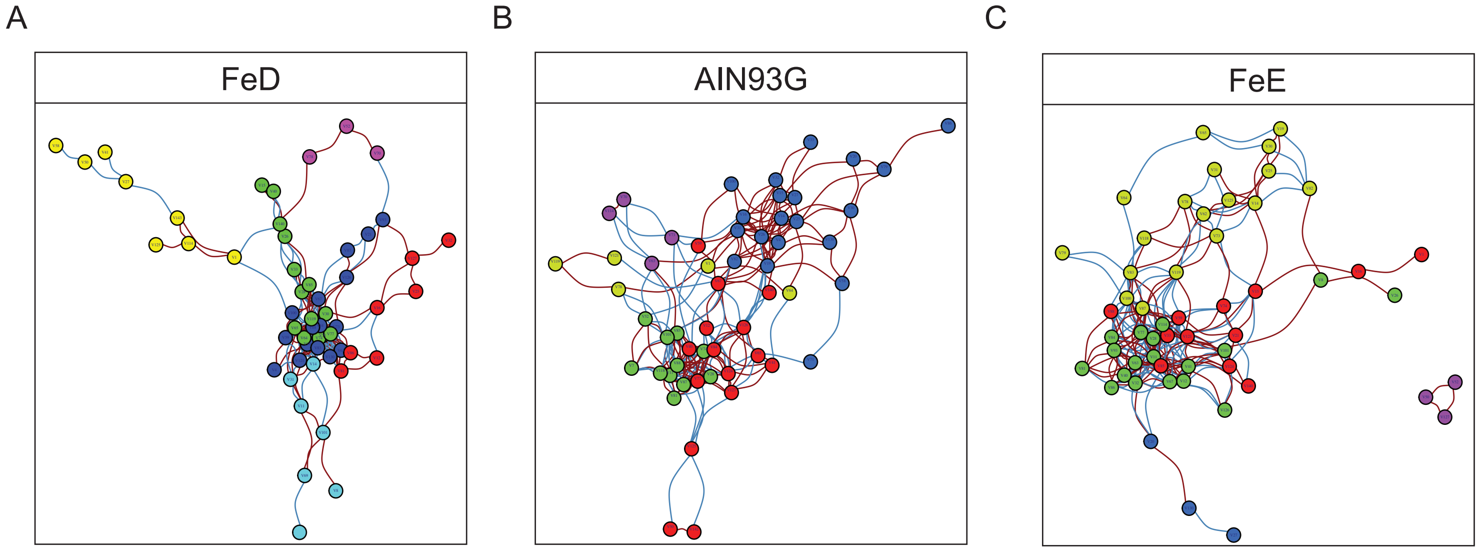

### Dietary iron content does not change the complexity of gut microbiota

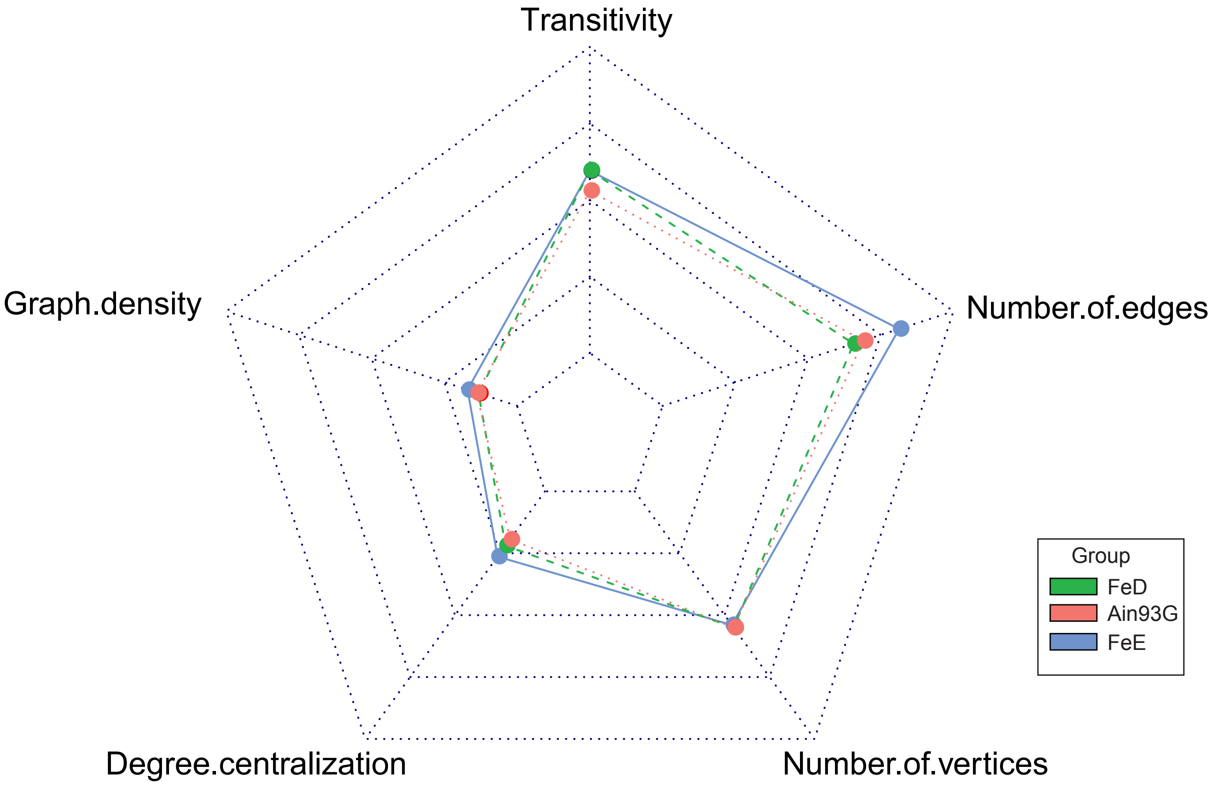

### Identification of the signature taxa by random forest

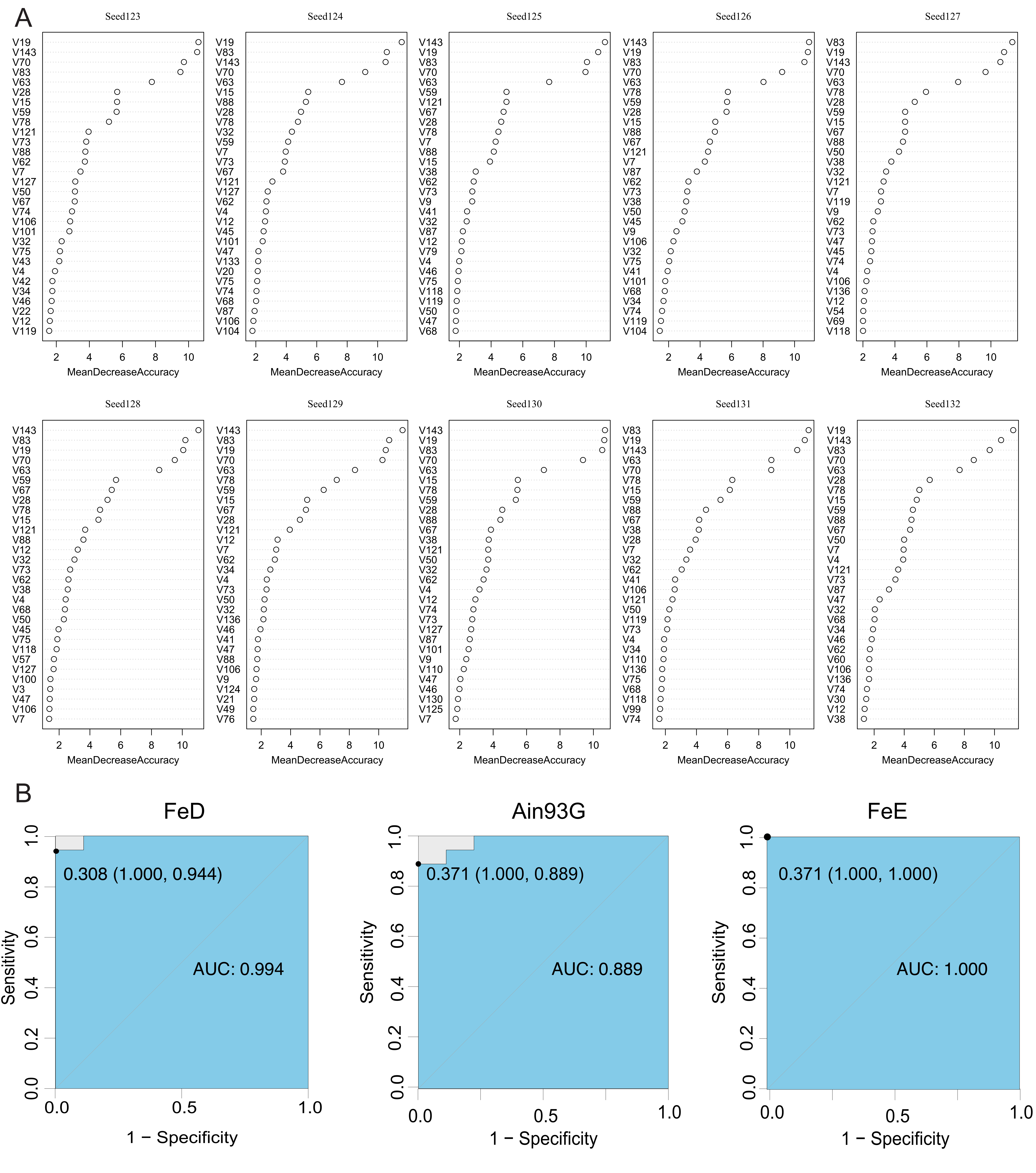

### Metagenomic prediction of microbial phenotypes using Bugbase

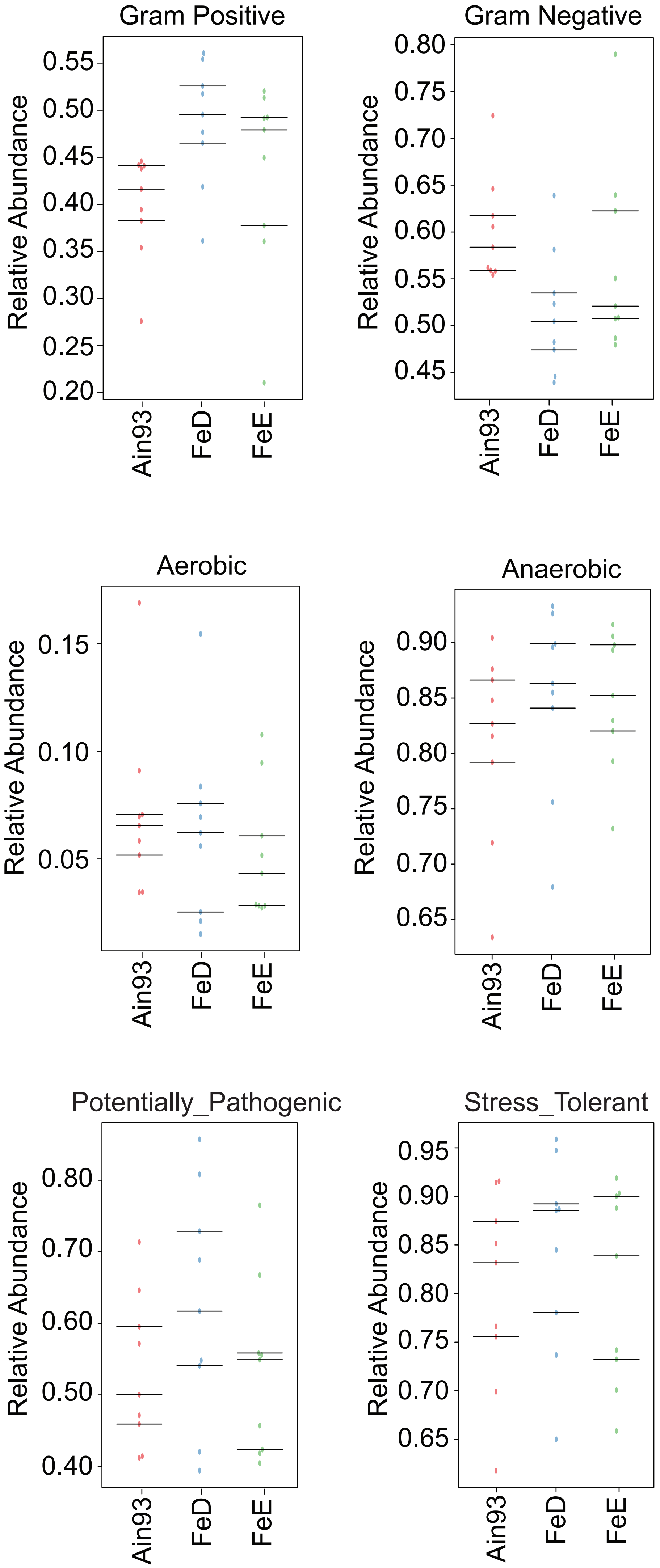

### Schematic representation of the link between gut microbiota and systemic iron homeostasis.

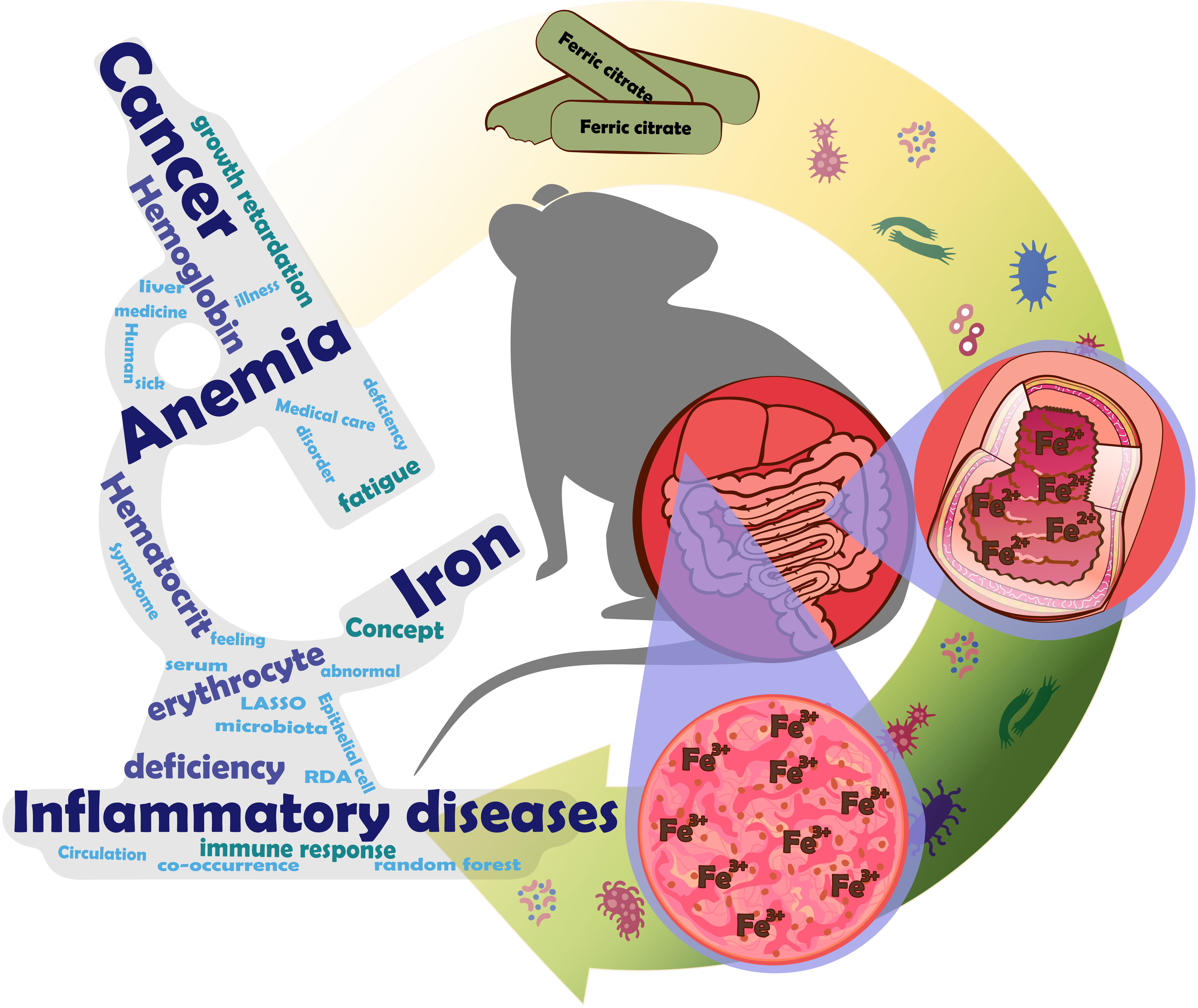

### Sequencing depth was assessed via the random sampling

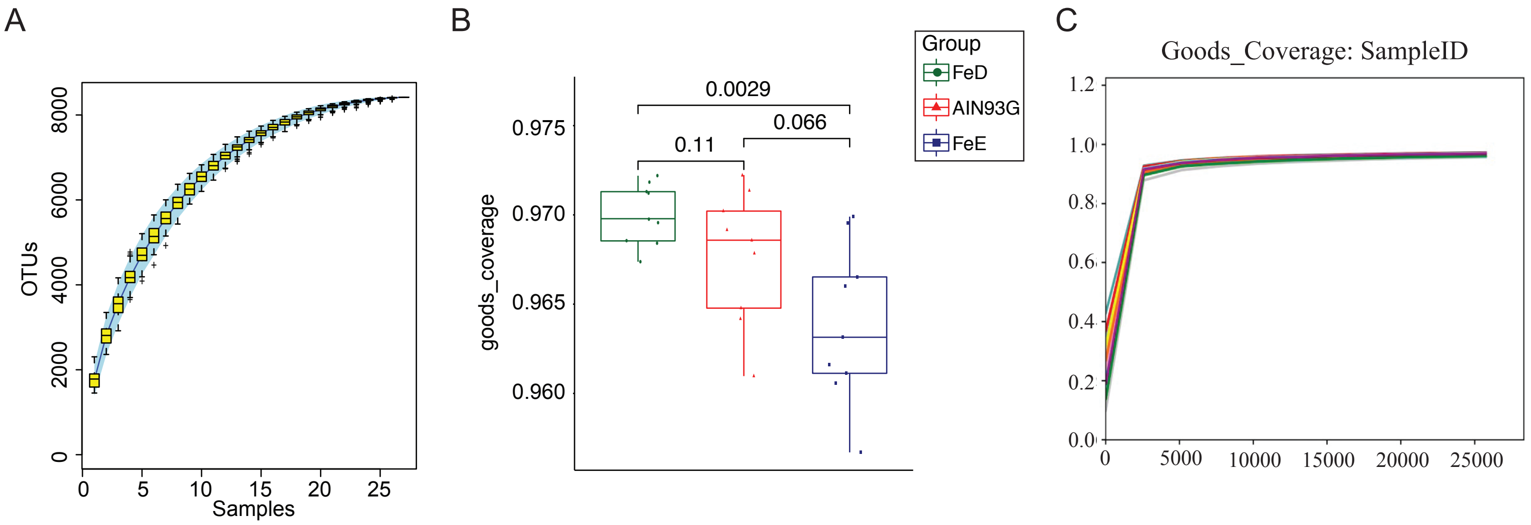
