## Supplementary material for "Fecal microbiota as a non-invasive biomarker to predict the tissue iron accumulation in intestine epithelial cells and liver": Identification of the signature taxa by random forest

A

| # | Bacteria |
|---|---|
| V19 | p__Bacteroidetes;c__Bacteroidia;o__Bacteroidales;f__Porphyromonadaceae;g__Parabacteroides;s__ |
| V83 | p__Firmicutes;c__Clostridia;o__Clostridiales;f__Peptostreptococcaceae;g__;s__ |
| V143 | p__Verrucomicrobia;c__Verrucomicrobiae;o__Verrucomicrobiales;f__Verrucomicrobiaceae;g__Akkermansia;s__muciniphila |
| V70 | p__Firmicutes;c__Clostridia;o__Clostridiales;f__Clostridiaceae;g__Clostridium;s__perfringens |
| V63 | p__Firmicutes;c__Clostridia;o__Clostridiales;f__;g__;s__ |

B

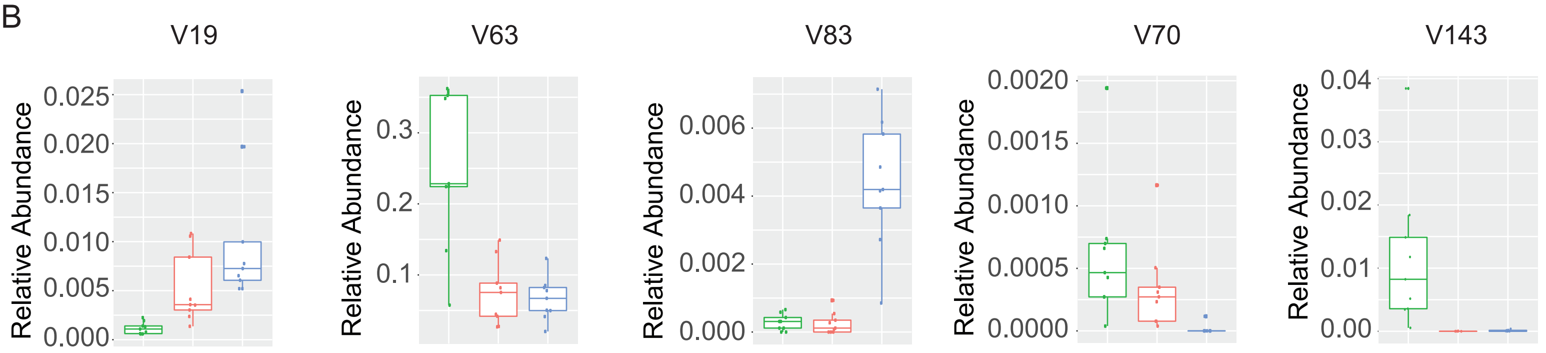


C

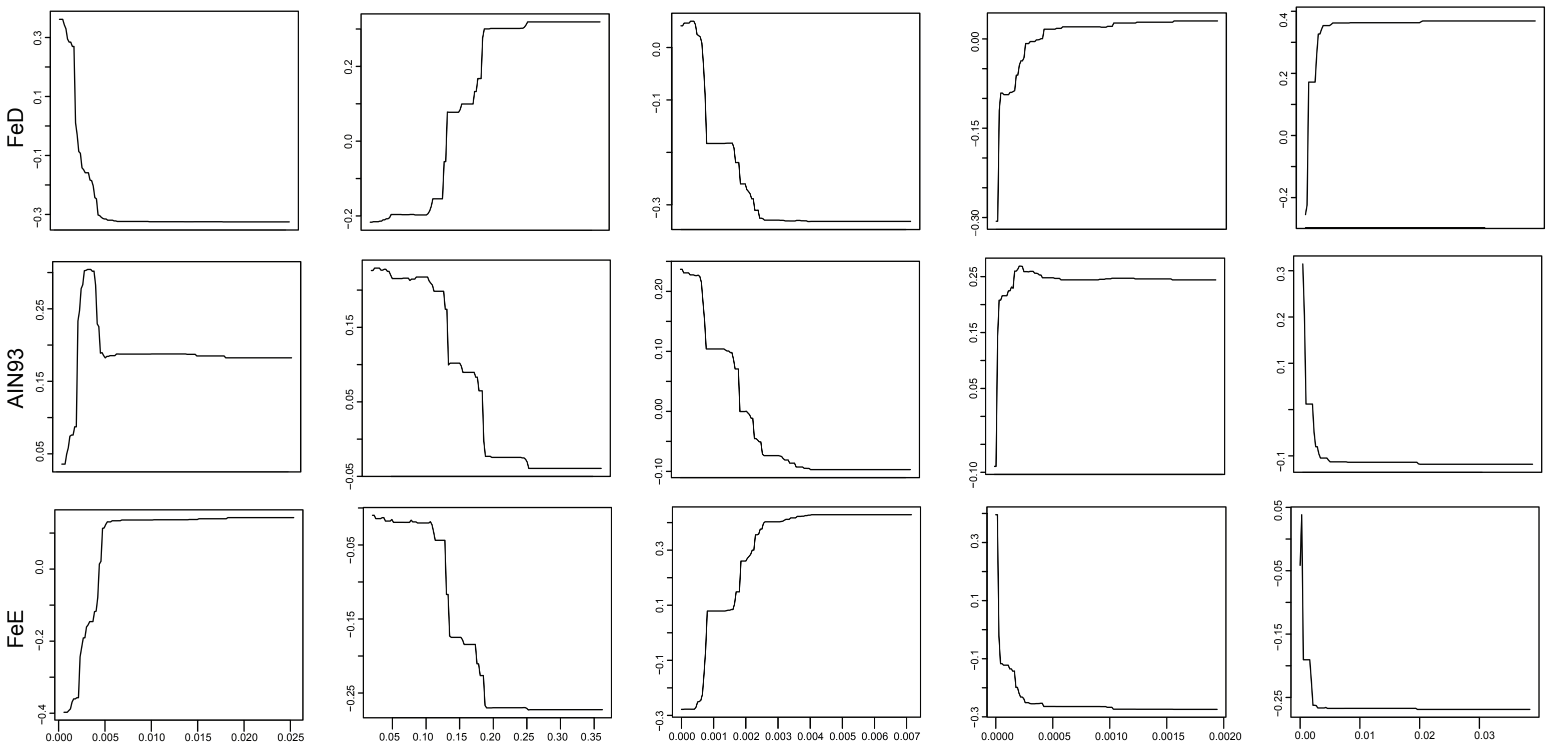


D

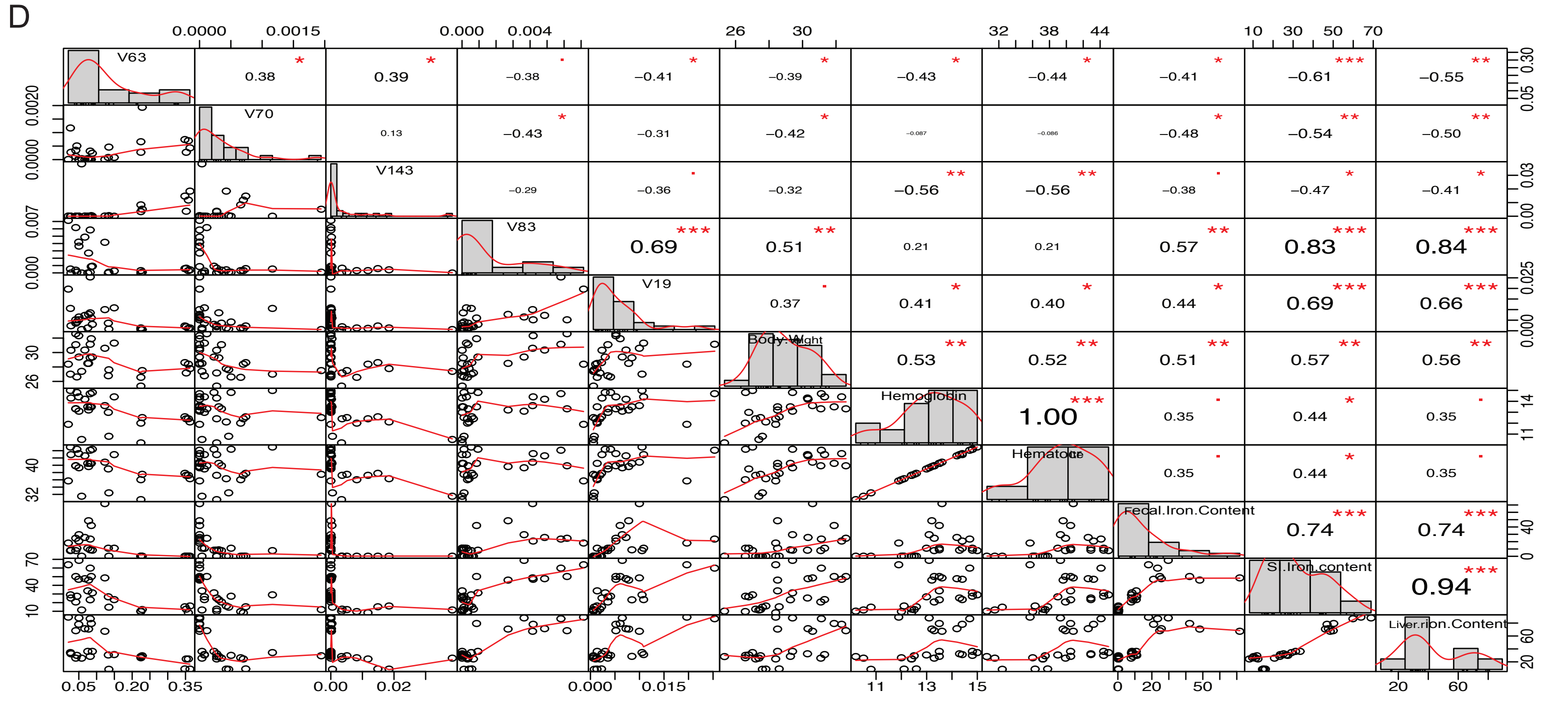
