## Supplementary material for "Fecal microbiota as a non-invasive biomarker to predict the tissue iron accumulation in intestine epithelial cells and liver": Selection of hyperparameter (???) for LASSO regression model.

cv_Fecal.Iron.Content lambda.min=3.46686940274949 lambda.1se=7.6449002856019

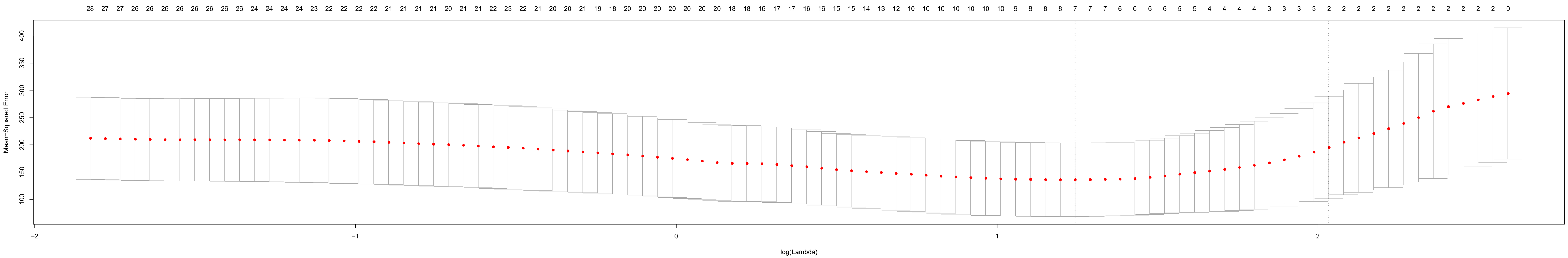


cv_SI.Iron.content lambda.min=0.226335150213356 lambda.1se=3.86433916911237

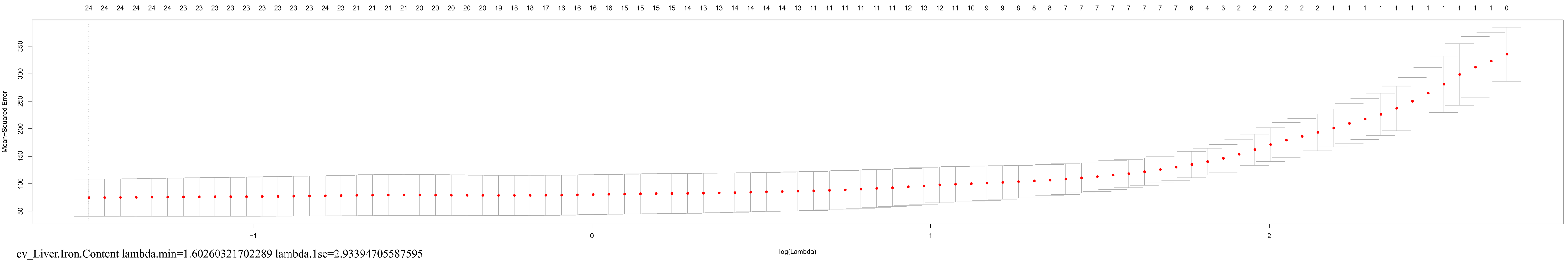


cv_Liver.Iron.Content lambda.min=1.60260321702289 lambda.1se=2.93394705587595

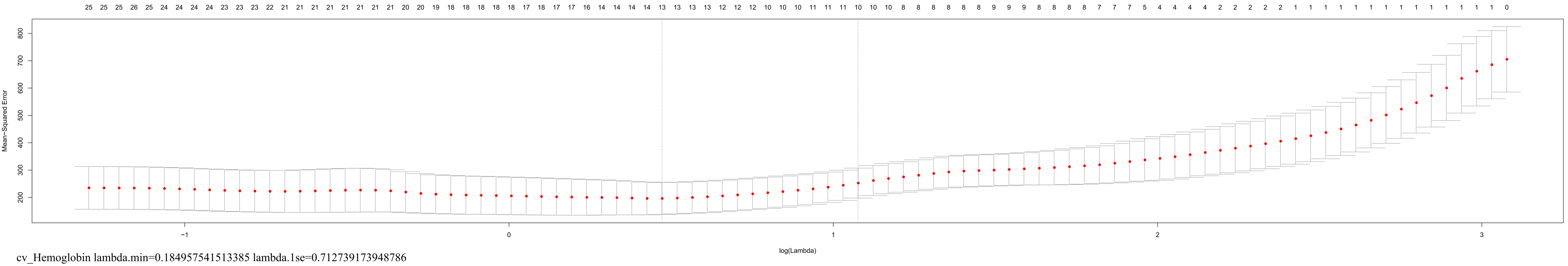


cv_Hemoglobin lambda.min=0.184957541513385 lambda.1se=0.712739173948786

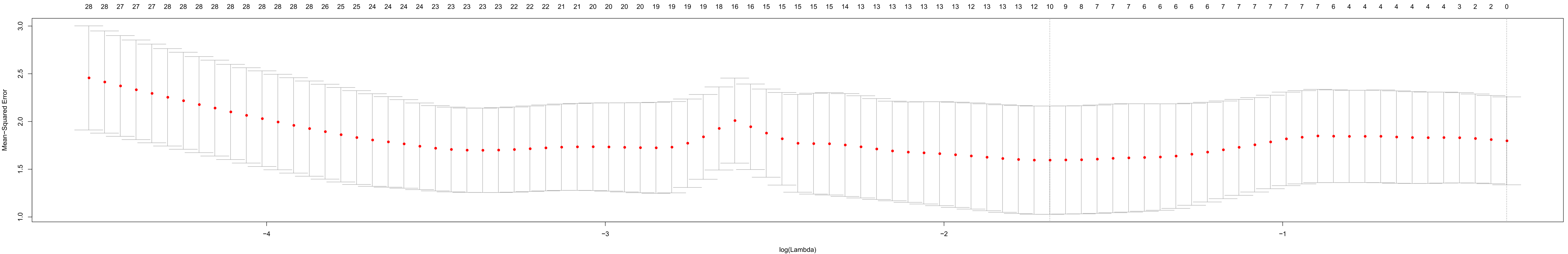
