## Supplementary material for "Fecal microbiota as a non-invasive biomarker to predict the tissue iron accumulation in intestine epithelial cells and liver": LASSO regression model to utilize the relative abundance of bacterial taxa to predict systemic iron levels

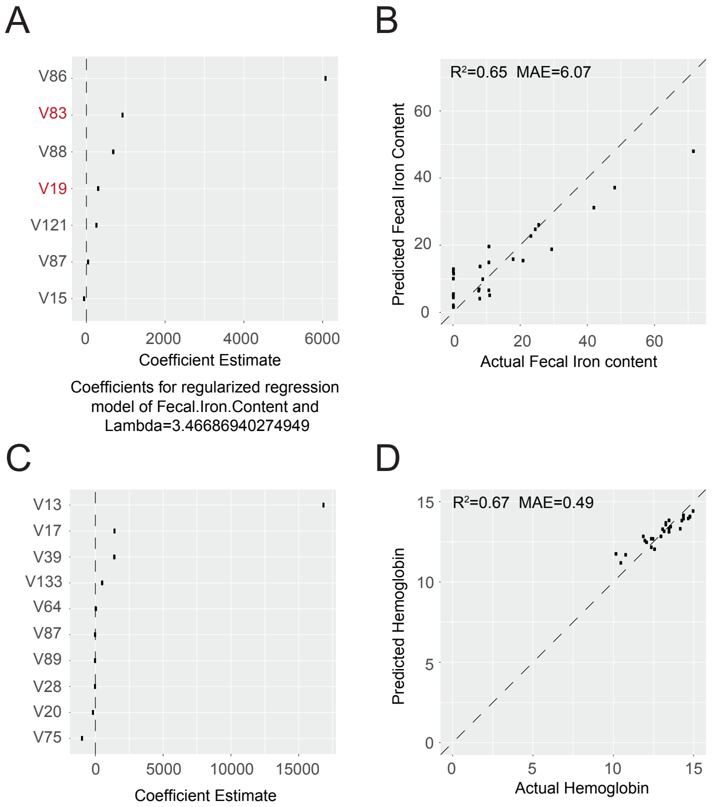


Coefficients for regularized regression model of Fecal.Iron.Content and Lambda=3.46686940274949

Coefficients for regularized regression model of Hemoglobin and Lambda=0.184957541513385
